# Can scientists fill the science journalism void? Online public engagement with science stories authored by scientists

**DOI:** 10.1101/760520

**Authors:** Yael Barel-Ben David, Erez S. Garty, Ayelet Baram-Tsabari

## Abstract

In many countries the public’s main source of information about science and technology is the mass media. Unfortunately, in recent years traditional journalism has experienced a collapse, and science journalism has been a major casualty. One potential remedy is to encourage scientists to write for news media about science. On these general news platforms, scientists’ stories would have to compete for attention with other news stories on hard (e.g. politics) and entertaining (e.g. celebrity news) topics written by professional writers. Do they stand a chance?

This study aimed to quantitatively characterize audience interactions as an indicator of interest in science news stories authored by early career scientists (henceforth ‘scientists’) trained to function as science reporters, as compared to news items written by reporters and published in the same news outlets.

To measure users’ behavior, we collected data on the number of clicks, likes, comments and average time spent on page. The sample was composed of 150 science items written by 50 scientists trained to contribute popular science stories in the Davidson Institute of Science Education reporters’ program and published on two major Israeli news websites - *Mako* and *Ynet* between July 2015 to January 2018. Each science item was paired with another item written by the website’s organic reporter, and published on the same channel as the science story (e.g., tourism, health) and the same close time. Overall significant differences were not found in the public’s engagement with the different items. Although, on one website there was a significant difference on two out of four engagement types, the second website did not have any difference, e.g., people did not click, like or comment more on items written by organic reporters than on the stories written by scientists. This creates an optimistic starting point for filling the science news void by scientists as science reporters.

## Rationale

The public draws primarily on the news media in general and internet news sites in particular for information about science and technology (1–4). Globally, digital media have supplanted traditional print and broadcast media, which has also affected science journalism (5,6). Today many science-related news items are written by part-time reporters or reporters specialized in other fields, who have less background and interest in covering science and technology (7–11). This shortage of specialized personnel has created an opening for the publication of public relations (PR)-generated content as journalistic content, which sometimes is even printed verbatim (12–17), thus relinquishing the traditional democratic role of the press as a watchdog that can signal misconduct, raise ethical questions and make critical observations.

A potential remedy for the declining numbers of professional science reporters was suggested in which scientists address the public directly (18–21). The argument is that by taking part in the public debate, scientists can contribute to influencing public discourse and policy (22). Visual scientists could counter fake news and constitute a role model for younger publics (23–25).

Correlational studies have shown that scientists who engage with the public also perform better academically (26,27). Web 2.0 provides scientists with platforms to directly disseminate their scientific messages, and allows broad audiences to comment, react, and potentially engage in dialogue with scientists (2,6,28,29). However, a closer examination of the audiences who interact with science on social media and dedicated blogs shows that they remain largely within the circles of academics and science enthusiasts (30,31). Hence, although social media platforms can increase the public’s exposure to science, the news media still wields distributional power that could be harnessed by scientists as a platform to present their ideas to wider audiences.

As noted in an editorial in ‘Nature’ in 2009: “An average citizen is unlikely to search the web for the Higgs boson or the proteasome if he or she doesn’t hear about it first on, say, a cable news channel. And as mass media sheds its scientific expertise, science’s mass-market presence will become harder to maintain”(19). Currently, scientists seldom have direct access to general news outlets. In addition, whereas scientists may be conversant in recent innovations and scientific breakthroughs, they are not skilled in writing in an engaging fashion for the public, particularly compared to media reporters.

Online news media adhere to different norms, agendas and styles than those found in the academic writing that scientists are accustomed to producing. The online news media compete for the public’s attention on a very tight schedule, that only allows a very short time for research, fact checking and the writing needed for science reporting, thus forcing journalists to operate under a heavy workload (15,32). While scientists write mostly for their peers to share, promote and advance scientific research, reporters aim to inform, alert and encourage public debate on topics that are thought to be on the public agenda or even purely entertaining (33–35). Whereas scientists are trained to write to other experts using a traditional, well accepted format of the IMRAD structure (Introduction, Methods, Results and Discussion) (20,36–38) and use scientific jargon abundantly, journalists use different genres and vocabulary to address non-expert audiences (20,36–40).

News sites are a competitive environment where scientists’ stories compete for attention with other news stories on hard (e.g. politics) and soft (e.g. celebrity news) topics (41) written by professional writers. Do they stand a chance? This study was designed to quantitatively characterize audience interactions with science news stories as an indicator of interest and attention. These stories, authored by early career scientists (henceforth ‘scientists’) trained to function as science reporters were compared to audience reactions to news items written by reporters and published in the same news outlets.

## Methodology

### Research context

The Davidson Institute, – the Educational Arm of the Weizmann Institute of Science in Israel has trained and employed graduate students, postdocs and research fellows in the sciences as writers for its website since 2006. In early 2014, an academic conference panel about science and risk communication in the online Israeli media^1^(The 6^th^ Israeli Science Communication Conference, (2015) Davidson Institute of Science Education, Weizmann Institute of Science, Rehovot (June 24-25)) hosted the editor in chief of the *Mako*^*2*^ (www.mako.co.il) news website. As a result of that meeting the Davidson Institute began collaborating with *Mako* by publishing science items written by scientists involved on its website (42). *Mako* is the third most visited Israeli News site (23.2M entries in the last quarter of 2016, (43)), which is owned and operated by ‘Keshet’, Israel’s largest TV commercial broadcasting company. It offers news content alongside streaming of TV shows. This type of collaboration was then also extended to *Ynet^3^*(http://www.ynet.co.il), Israel’s most widely read news website (52.5M entries in the last quarter of 2016, (43)). *Ynet* is operated by the ‘Yediot Aharonot’ media group that publishes a daily tabloid newspaper in addition to the website and caters mostly to young audiences (aged 18-34) surfing on mobile devices (43). Both news sites provide freely available news content. The two websites do not employ a dedicated science journalist, or require the reporters covering these topics to have background in science.

Currently, the Davidson reporters program employs about 50 graduate science students and faculty who attend an annual brief training program led by the editorial team that focuses on practical methods to improve popular writing (e.g., avoiding jargon and passive voice). The writing process is mostly based on individual contacts between the scientists and the in-house editor. The Davidson editorial team is composed of science editors, two content editors and an editor in chief who is a former journalist. All the editors, except one content editor, have academic science degrees. In cases where the content editors have reservations about the content, they consult the scientist who authored the item before sending it for a final revision by the editor in chief. The scientists have backgrounds in different fields and are at various stages of their graduate degrees, although a few are already faculty members. There is no quota demanded of each writer, but most scientists write between one to four 500-word items a month. The topics span the science, technology, mathematics and engineering (STEM) fields, and are chosen by the editorial team as a function of their newsworthiness and potential for public interest (or based on topics suggested by the scientists). The scientists are not allowed to write about their own research or research done in their labs, but are encouraged to write about local Israeli research as part of the Davidson agenda. The scientists are employed on an hourly basis to promote thorough inquiry (rather than being paid on the basis of number of words).

The Davidson Institute initially proposed this collaboration with the *Mako* and *Ynet* news site editors to increase the quantity and quality of science content in the news. According to Davidson staff, this collaboration enables scientists to share accurate, innovative scientific information and make it part of public’s everyday news consumption while the news sites benefit from free high-quality science content. To date, this arrangement involves most of the mainstream news sites operating in Hebrew in Israel. The news site editors are provided with edited text, which they are not allowed to alter without Davidson’s permission, but they are free to change the headlines.(For a critical analysis see also (42)). The name of the scientist appears in the credit line, and is visible to the readers even before clicking the item to read, and includes the person’s title and affiliation to the Davidson Institute. The name of the writer, his or her affiliation and status is also stated at the end of the item (e.g. “Yael Groper, Davidson Institute of Science Education website reporter and a doctoral student in the Weizmann Institute of Science”) alongside a link to the research article, if the item is based on one.

#### Researcher positioning

The first and third authors are university-based science communication researchers not affiliated with either Davidson, *Mako* or *Ynet*. The second author is the head of the science communication unit in the Davidson Institute and the initiator of the scientist writing program and collaboration with the news sites.

### Data source and sampling

Digital media and Web 2.0 allow access to accurate and updated data on consumers’ engagement with content (44,45), and sometimes even influence editorial decisions on topics and item placement (46–48). Previous studies have pointed to the disparity between what journalists think interests the public and readers’ choices, mainly as regards the emphasis on public affairs issues (49). Studies of public engagement with science content online have mainly focused on views (clicks) and comments. They have analyzed engagement in terms of different forms of interactivity offered by the online platforms (e.g. clicking, commenting, emailing, etc.) that varies between topics and is time and context sensitive (44,50–54).

Although engagement data are used routinely in online newsroom decision making, little is known about the characteristics of public interactions with online science items (55). This is due primarily to the difficulty of obtaining data (e.g., number of clicks and average time on page) which are kept confidential for commercial reasons.

As part of this research-practice collaboration, the researchers were given access to confidential Google Analytics data, including clicks, time spent on the page, likes and comments on *Mako’s* and *Ynet*’s websites. Due to the commercial sensitivity of the information, data mining took place on several consecutive days at the news company offices at *Mako*. The researcher was not allowed access to the information directly from *Ynet* but was sent the requested data electronically. Data were kept on an encrypted drive with access restricted to the first author alone. Due to the commercial nature of the data, the researchers were committed not to disclose the raw data. Hence, here, only descriptive data such as averages and standard deviation are presented.

The initial dataset was composed of all 296 news items written by scientists and published on the two websites from July 2015 to January 2018 (Table 1). These were published mainly on the Digital, Health, Animals, Culture and Traveling channels (ranked by the number of items in each section).

**Table 1.** Data collection. Data were collected in March 2016 and January 2018. Dark grey shading indicates months when scientists’ items were published on the *Mako* website (n = 114) and in light grey for *Ynet* (n = 182). The matching process yielded a total of 150 pairs of news items. Items that did not have a corresponding item written by reporters (n=57) and items without access to the full engagement data (n=89) were omitted.

| Year | Jan |  | Feb |  | Mar |  | April |  | May |  | Jun |  | Jul |  | Aug |  | Sep |  | Oct |  | Nov |  | Dec |  |
| --- | --- | --- | --- | --- | --- | --- | --- | --- | --- | --- | --- | --- | --- | --- | --- | --- | --- | --- | --- | --- | --- | --- | --- | --- |
| 2015 |  |  |  |  |  |  |  |  |  |  |  |  | 3 |  | 2 |  | 1 |  | 6 |  | 6 |  | 7 |  |
| 2016 | 6 | 1 | 4 | 1 | 10 | 4 | 6 | 2 | 3 | 2 | 4 | 4 | 4 | 9 | 6 | 10 | 0 | 0 | 0 |  | 8 | 13 | 5 | 15 |
| 2017 | 2 | 14 | 2 | 11 | 5 | 15 | 7 | 13 | 5 | 18 | 4 | 14 | 4 | 16 | 4 | 10 | 7 |  | 0 |  | 1 |  | 1 |  |
| 2018 | 1 |  |  |  |  |  |  |  |  |  |  |  |  |  |  |  |  |  |  |  |  |  |  |  |

### Database of matching items

Each science item written by a scientist was paired with a corresponding news item written by a professional reporter that was published on the same channel and within an average of 0.8 days apart (about half of the items were published on the same day, and the rest no more than three days before or after); see Figure 1 for an example of paired items. After this initial filtering by channel and date of publication, we made efforts to pair similar formats (e.g. quizzes, video articles, short/long items, etc.) when there was a choice of several organic items. In cases where no corresponding item was found on the same channel and within the designated timeframe, the reference item was excluded from the database (n = 57). Due to restricted access of the researcher to the *Ynet* news site’s data and to broken links in both news sites, another 89 pairs were omitted from the sample. Overall, the process yielded a total of 150 pairs of news items, 51% of the initial dataset. The final database is a representative sample of the full database consisting of all channels and within the same time range as the full database.

**Figure 1.**
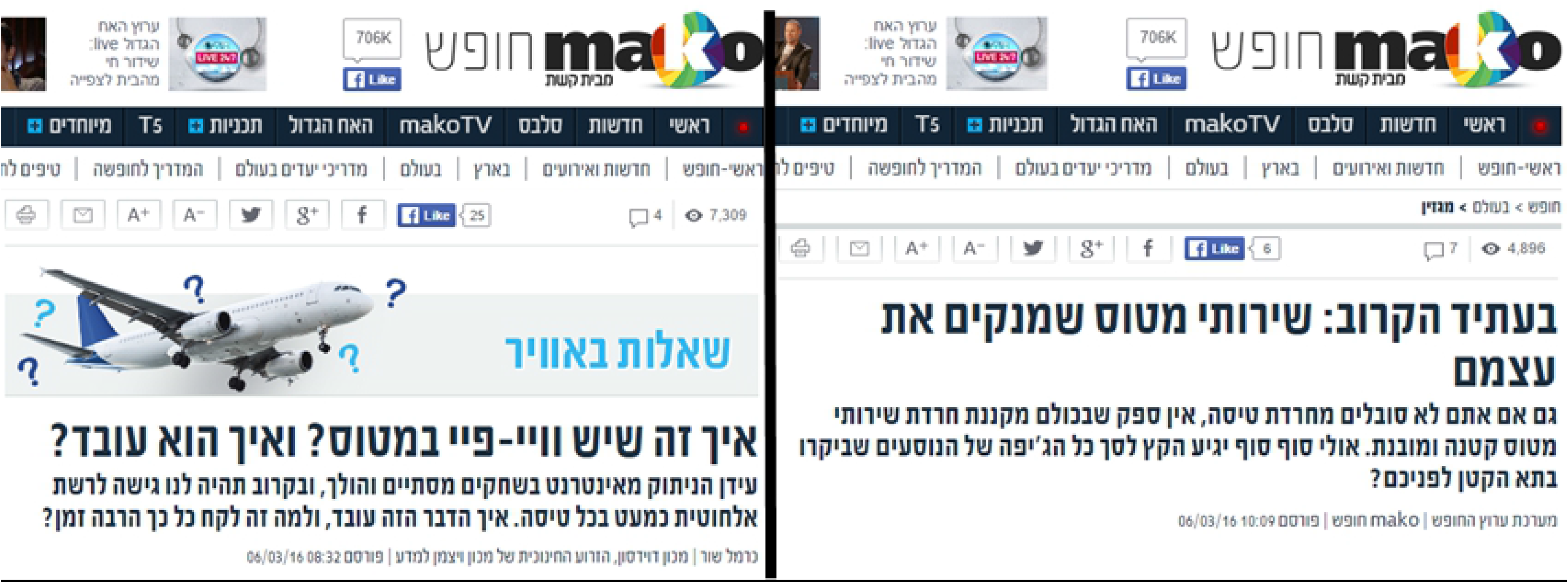
Screenshot of paired items from the *Mako* website to illustrate the matching process. The item on the left, written by a scientist is titled “How can we have Wi-Fi on an airplane? And how does it work?“(written by Carmel Shor), whereas the item on the right was written by the website’s reporter and was titled “In the near future: airplane toilets that clean themselves” (written by the vacation channel editorial). Both items were published on the same day and on the same website channel (“Vacation”).

On *Mako* website, 69% of items written by scientists were published on the Health channel, alongside site reporters’ items on new treatments and diet suggestions (e.g. “Five delicious recipes for a healthy meal”, 21/8/2017). Another 12% of the scientists’ generated items were published on the Holiday/Travel channel and paired with items on new popular vacation resorts, celebrities’ vacations and other travel information (e.g. “Where do the residents of the “Big Brother” reality show love to go on vacation?”, 1/1/2016). An additional 7% were published on the HIX magazine devoted to “Scientific discoveries, interesting phenomena, funny inventions, exciting news and other events from the world”, and on the Culture channel.

On *Ynet* website, the vast majority of items (97%) were published on the Digital channel, alongside items such as “After a mouse and a gorilla: was a shark photographed on Mars?” (27/3/2016). The remaining 3% were published on the Animals, Health and World channels. While the scientists’ items were always about a scientific study, or science related issues, the paired organic items were more diverse in terms of topics. Although the paired items had a similar topic, since they were published on the same channel, an organic item on the Digital channel on *Ynet* for example, could be a set of pictures showing readers what the Earth looks like at night, without any scientific or research related content. In four rare occasions both paired items covered the same exact topic side by side (one on *Mako* and 3 on *Ynet*, see Table 2 for item headlines). About half of the pairs in the database were both scientist and organic reporter items, on a scientific or research related topic. Since we could not pool the paired items from the two websites, there are too few items from each site to calculate statistical significance.

**Table 2.** Headlines of paired items on the same topics. Four rare occasions in which an item on the same topic was published written by the website’s organic reporter and also by a scientist from the reporters’ program. Items addressed the same topics and were published more or less at the same time. These items differed in terms of the frames and angles the writers chose to take. Public engagement was higher with the items written by reporters on the first and last items, whereas the opposite was found for the two others.

| Website | Scientist's item headline | Reporter's item headline |
| --- | --- | --- |
| *Ynet* | “Seeing through Jupiter’s clouds”<br>(4/7/2016) | “After a 5-year journey: Juno will reach Jupiter<br>this week” (3/7/2016) |
|  | "An Earth-like planet was discovered 4 light years away" (24/8/2016) | "Has a planet similar to Earth been found?" (24/8/2016) |
|  | "A solar system with 7 Earth-like planets was discovered" (22/2/2017) | "Seven planets right next to each other: "Not much chance for life"" (22/2/2017) |
| *Mako* | "The HPV vaccination is perfectly safe" (21/10/2016) | ""The vaccination doesn't cause paralysis that same day"" (20/10/2016) |

### Data analysis and engagement types

Four quantitative parameters were chosen as indicators of audience engagement based on previous studies (43–45,51–55) and the available data. The data used in this study relied solely on absolute numbers for each parameter. Other relevant variables, such as an item’s location on the website, length of time visible on the channel, length of the item etc., were not available. ***Clicks*** *(views)* were used as an indicator of interest in reading the item based on the headline. When presented on the entry page, a secondary sub-headline was visible to readers as well. The number of clicks ranged widely from a low of 651 to a high of 269,802 clicks on the most popular item “How does Bitcoin work?” (*Ynet*, Dec. 2017) (for average clicks per site see Table 3).

**Table 3.**
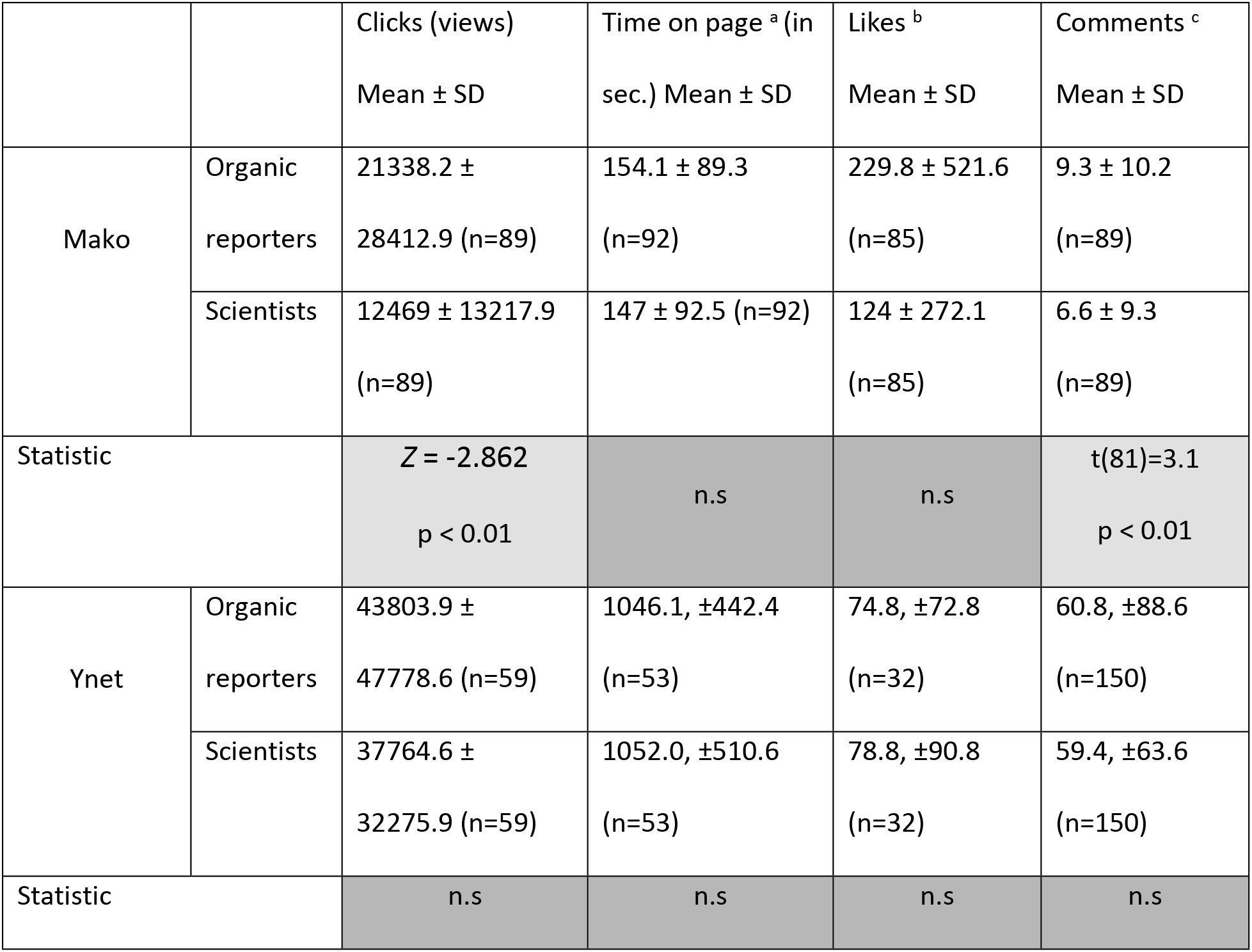
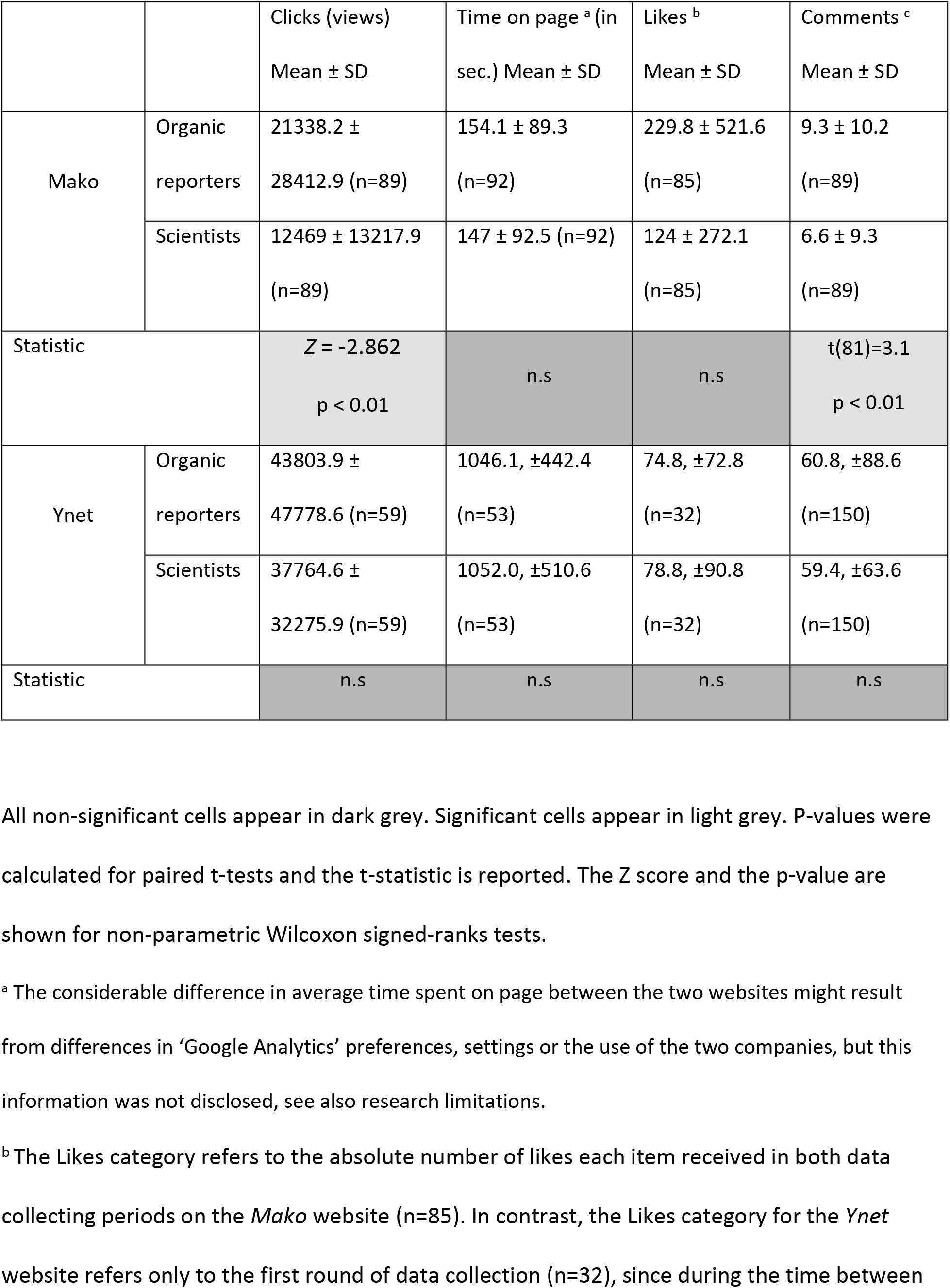
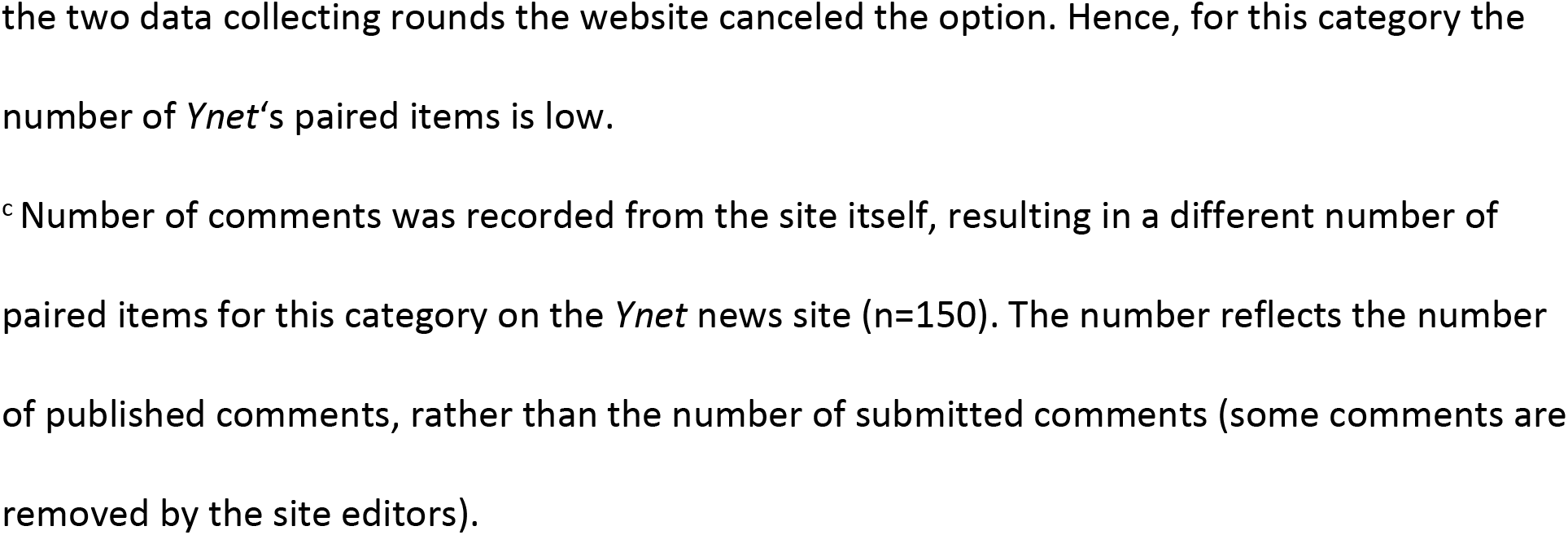
Comparison of reader engagement with items by scientists and reporters.

***Average Time on page*** (reported in seconds) represents the time devoted by readers to each item. This indicator can provide some indications as to whether the item was read in full. The average time spent on page ranged from 13 seconds to 1,702 seconds for the item “The God Pan and nude festivals in the Golan heights 1900 years ago” written by the website’s reporter (*Mako*, June 2016) (for averages per site see Table 3).

***Likes*** can represent readers’ favorable opinion of the item or the event it describes. It demands a higher engagement level on the part of the reader since by clicking ‘Like’ (in the Mako website, the ‘Like’ option is marked as ‘Recommend’), the item is published on the reader’s wall on Facebook, thus exposing it publicly. At the time of data collection *Facebook* was the only social media platform with an available interface with the two news sites. Likes ranged from zero to 3,600 for the most popular item “An Earth-like planet was discovered 4 light years away” (*Ynet*, August 2016) (for averages per site see Table 3).

***Comments*** require more time and effort relative to ‘Likes’. Comments could be one word long to several paragraphs long, and may be off topic. Any internet user can post a comment anonymously on these two websites. The number of comments ranged from no comments at all to 621 comments on the most popular item “The physics of building pyramids” (*Ynet*, April 2017) (for averages per site see Table 3).

### Statistical analysis

Each engagement type was assessed for normal distribution. A paired sample t-test was used for normally distributing parameters, such as average time on page on both websites and comments on *Mako* and Likes on *Ynet*. A Wilcoxon signed-rank test was used for parameters that were not normally distributed, such as Clicks on both websites and Likes on *Mako* and comments on *Ynet* (α=0.05). In order to verify that the time difference between the two sampling cycles did not affect the data, a one-way ANOVA was run on the first and last two months and the median date of the data collection cycles for each parameter. No significant difference was found. Therefore, the results of the statistical analyses are presented for the two rounds of data collection combined.

## Results

In this study we are in the unique position where non-significant differences between groups are highly informative for the goals of the study. Figure 2 presents the comparison of all engagement types on both websites.

**Figure 2.**
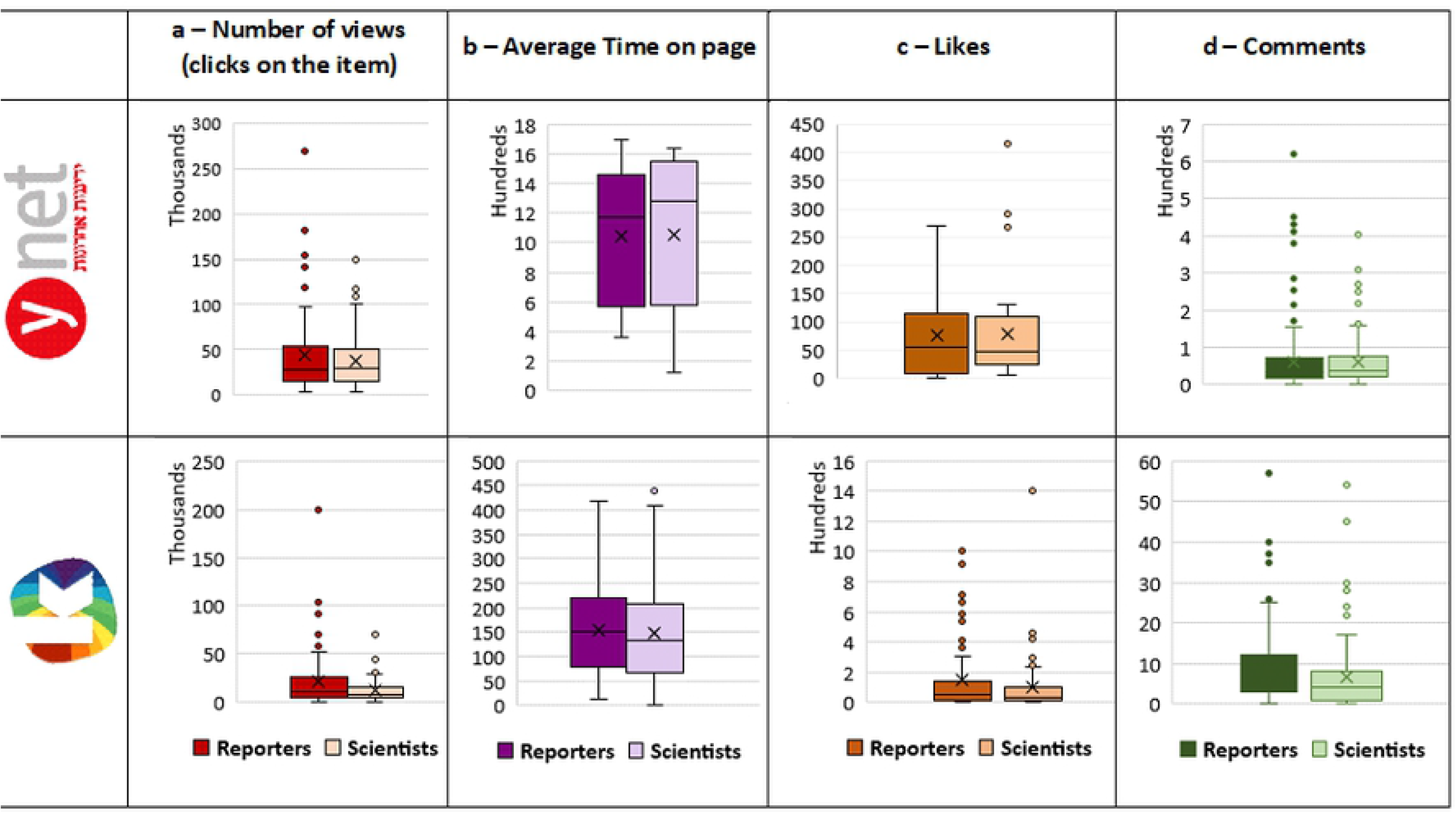
Number of views, average time on page, likes and comments in items authored by reporters and items written by trained scientists. Top row portrays data from the *Ynet* news site, lower row shows data from the *Mako* news site. Column a. distribution of number of Clicks on items (*Ynet* n=59 pairs, *Mako* n=89 pairs); column b. distribution of average time spent on page (in sec.) (*Ynet* n=53 pairs, *Mako* n=92 pairs); c. distribution of the number of likes received (*Ynet* n=32 pairs, *Mako* n=85 pairs); column d. distribution of the number of comments on items written by reporters vs. trained scientists (*Ynet* n=150 pairs, *Mako* n=89 pairs).

In the case of the *Mako* website no significant differences were found between items written by scientists and *Mako*’s organic reporters for average time on page and ‘Likes’, based on 92 and 85 pairs of items, respectively (Table 3). On the other hand, a Wilcoxon signed-ranks test showed a statistically significant difference in the number of Clicks (views) between items written by scientists and organic reporters (n=89, *Z* = −2.862, *p* = 0.004) and a paired sample t-test showed a statistically significant difference in the number of Comments (n= 89, p<0.05) with more public engagement in response to *Mako*‘s’ organic reporters on both parameters. To conclude *Mako*’s results, there was no difference in the time devoted to reading the items or liking them (hitting the ‘Like’ button) but there was a clear preference to click items that were not written by scientists and were not necessarily about science.

While *Mako*’s results were mixed, on the *Ynet* news site an analysis of the data retrieved on the paired items showed no significant differences between items written by scientists and organic reporters on any of the parameters (Table 3). For example, the average length of time on page for items written by organic reporters was 17:26 minutes, whereas the time on page for an item written by scientists had a viewing duration of 17:32 minutes, on average, as shown in Figure 1. Thus, based on our data, the public’s interactions with science news written by scientists were not significantly different from other news items written by reporters on *Ynet’s* news site.

## Discussion

Americans’ and Israelis’ primary information source about science and technology is the online news medium (4,56), which is impacting public attitudes, perceptions and even behavior (1,57–61). News media has the potential to promote scientific understanding, especially if sufficient explanations of the science are provided and a narrative is used for its presentation (57).

Hence, accurate, accessible and relevant science stories in the news media are important factors in a healthy science communication environment. Unfortunately, given the current collapse of traditional journalism worldwide, science journalism has become a major casualty, thus hindering its ability to provide quality science coverage. For example, between 2013 and 2015, the number of science reporters in the Israeli news media decreased from 9 to only three, a drop that continues to this day (62). The results of this study suggest that filling the void created by the firing of titular science reporters by scientists who write for the media may be a viable solution resulting in a higher frequency of scientific content in the news media which is also attractive to readers.

This study examined whether readers reacted differently to science news items written by scientists as compared to news items written by organic reporters published on the same online news media sites. Generally speaking, based on our findings, the answer is no: audiences interacted similarly with both. This finding justifies the time and effort invested by the scientists and the Davidson science communication team to write attractive science stories, and justifies the resources provided by the news sites. Apparently if websites publish it, audiences will consume it. The website gains highly credible, reliable, science items free of charge and the scientists get a spot in a high exposure platform (63).

These optimistic results raise a normative question about the impact of these collaborations on science journalism and science communication. In a democratic society the media do not only serve as an information conduit for the public (64), but also as platform to critique the authorities as regards misconduct, corruption or misuse of public resources (17,64). Scientists communicating science while working as scientists can fill the informing and popularizing void, but cannot take on the watchdog role since they cannot be expected to provide a critical independent outsider approach of journalists. Providing ready-made scientific content to news outlets without charge may perpetuate the media’s reliance on free, external content, and hence contribute to the weakening position of science journalists. It could be argued that accustoming the media to getting ready-made content without journalistic scrutiny may in fact be advancing ‘churnalism’- a practice in which pre-packaged stories provided by news agencies and press releases are adapted for publication instead of reported news, and therefore potentially posing a danger to the legitimacy of science journalism and undermining its credibility (12,13,65).

While these are important caveats, it is crucial to note that although the Israeli online news media form one of the most accessible sources of information about science and technology related topics, they are also ranked as mostly unreliable (an average of 5.6 on a 10-point scale). University scientists, on the other hand, are seen as the most reliable source of information (an average of 7.6 on a 10-point scale), but inaccessible (only 19% of the survey respondents mentioned them as their sources of information about science and technology) (66). The model described here integrates the strengths of each source; i.e., exposure and accessibility on online media and the reliability and expertise of university scientists.

The results from the two sites were mixed. No significant differences were found between scientists’ and reporters’ items on *Ynet*, but mixed findings were found on *Mako*, where reporters’ items had more views and comments, but likes and time on page did not differ. The differences between the results from the two news sites may be attributed to their characteristics: *Mako* is owned by ‘Keshet’, Israel’s largest commercial TV broadcasting company and shares the same website (www.mako.co.il) with television content on demand.

*Ynet* is operated by the ‘Yediot Aharonot’ media group that also publishes a daily newspaper. This may suggest differences in the expectations and motivations of audiences reading the two websites. Although both websites offer news coverage on an array of topics, most visitors to the *Mako* website expect more entertaining options than when accessing *Ynet* for more hard news content. Given these differences in press orientation, our assumption is that more *Mako* readers accessed the website with light entertainment in mind, rather than an interest in finding out about recent scientific or technological developments. Another explanation is related to the structure of the websites. While *Ynet* has a designated channel for science in which the majority of the items in the database were published (97% of the items on *Ynet*, 39% of the total database), *Mako* does not publish this type of channel, and all the items from the database were published in the different channels the website offers its audience.

This study was enabled by a Research-Practice-Partnership (RPP). This unique position allowed the researchers access to data that are usually unavailable for commercial reasons, while providing practitioners with an evidence-based evaluation of their science communication efforts and the public’s interaction with their products. Such RPPs have immense potential for improving practice and tapping experience-based questions and real-life data. More of these collaborations will increase reliance on behavioral data that can complement self-report research instruments.

### Research Limitations

One of the key limitations of this study was its reliance on the *Google Analytics* data mining system. The algorithm used to collect data by *Google* is unknown, and so are the basic assumptions underlying its data mining algorithm. Hence this study relied solely on absolute data for the number of clicks (views), Likes, etc. We did not use other information provided by *Google Analytics* such as age and gender since these constitute inferable data that rely on *Google’s* search and deduction algorithm.

Our data show a significant difference in average time spent on page between the two websites. This could also be a result of the settings, preferences and specifications each news company used to configure the data collection. These were not disclosed either. This problem was mitigated by not comparing public engagement between the two websites, only between writers on the same website.

Finally, it is important to emphasize that this study focused on the quantity and type of audience interactions with two types of coverage. The quality of the coverage and user generated content was not addressed. For example, comments might only contain a title with no additional text, or be several paragraphs long. Comments could be positive or critical (e.g., ‘interesting but it was hard to understand’) or off topic. The quality of these aspects might defer between the types of coverage and their associated reader comments.

## Conclusion

This study examined the public’s interactions with scientists’ popular writing, thus shedding light on online public engagement with science outside of an experimental setting. The results paint a positive picture where in most cases no differences were found between the ways audiences responded to scientific reports written by scientists-as-science-reporters, and stories written by news site reporters. This model may thus provide a practical solution for filling the science journalism void on struggling news media.

## Acknowledgments

Removed for review

